# The demographic drivers of cultural evolution in bird song: a multilevel study

**DOI:** 10.1101/2023.11.30.569452

**Authors:** Nilo Merino Recalde, Andrea Estandía, Sara C. Keen, Ella F. Cole, Ben C. Sheldon

**Affiliations:** Edward Grey Institute, Department of Biology, University of Oxford, Oxford, UK; Earth Species Project, 1536 Oxford St. Berkeley CA 94709, US

**Keywords:** animal culture, bird song, demography, cultural evolution

## Abstract

Social learning within communities sometimes leads to behavioural patterns that persist over time, which we know as culture. Examples of culture include learned bird and whale songs, cetacean feeding techniques, and avian and mammalian migratory routes. Shaped by neutral and selective forces, animal cultures evolve dynamically and lead to cultural traditions that differ greatly in their diversity and stability. These cultural traits can influence individual and group survival, population structure, and even inform conservation efforts, underscoring the importance of understanding how other population processes interact with social learning to shape culture. Although the impact of social learning mechanisms and biases has been extensively explored, the role of demographic factors—such as population turnover, immigration, and age structure—on cultural evolution has received theoretical attention but rarely been subject to empirical investigation in natural populations. Doing so requires very complete trait sampling and detailed individual life history data, which are hard to acquire in combination. To this end, we built a multi-generational dataset containing over 109,000 songs from >400 individuals from a population of Great Tits (*Parus major*), which we study using a deep metric learning model to re-identify individuals and quantify song similarity. We show that demographic variation at the small spatial scales at which learning takes place has the potential to strongly impact the pace and outcome of animal cultural evolution. For example, age distributions skewed towards older individuals are associated with slower cultural change and increased diversity, while higher local population turnover leads to elevated rates of cultural change. Our analyses support theoretical expectations for a key role of demographic processes resulting from individual behaviour in determining cultural evolution, and emphasize that these processes interact with species-specific factors such as the timing of song acquisition. Implications extend to large-scale cultural dynamics and the formation of dialects or traditions.

## RESULTS AND DISCUSSION

Some behavioural traits are shared and persist within communities due to social learning (Viciana, 2021). We refer to these behaviours as ‘animal culture’, exemplified by tool use in capuchin monkeys (Falótico et al., 2019), the learned songs of oscine birds, migration routes (Berdahl et al., 2018; Byholm et al., 2022; Jesmer et al., 2018), and the feeding techniques of some cetaceans (Allen et al., 2013; Rendell & Whitehead, 2001). Animal cultures are not static: neutral and selective mechanisms influence the frequency of cultural traits (Potvin & Clegg, 2015; Williams & Lachlan, 2021), leading to a process of cultural evolution. The resulting cultural traditions differ considerably in their diversity and stability (Tchernichovski et al., 2017), determined by both learning biases and mechanisms and the demographic structure of populations (Deffner et al., 2022; Kandler et al., 2017).

While the role of social learning strategies and biases—frequency dependence, tutor biases, etc.— has been extensively studied (Aplin et al., 2017; Kendal et al., 2015; Lachlan et al., 2018; Pike & Laland, 2010; Tchernichovski et al., 2021), there exists a substantial gap in our understanding of how demography contributes to the emergence and persistence of distinct cultural traits within wild populations. Processes such as the recruitment of juveniles and immigration, emigration, mortality, and variation in age structure are likely to strongly affect an individual’s opportunities for learning and exposure to different cultural variants, which has been amply emphasised by theoretical work (Barta et al., 2023; Deffner & McElreath, 2020, 2022; Deffner et al., 2022; Derex & Boyd, 2016; Fogarty et al., 2019; Kandler et al., 2023; Kirby & Tamariz, 2021; Nunn et al., 2009). Despite this, translating theoretical expectations into empirical evidence remains a challenge (see Chimento et al., 2021; Fayet et al., 2014 for exceptions).

Culture is increasingly recognised as both a fundamental aspect of many animals’ lives and a valuable tool in monitoring and conservation efforts (Brakes et al., 2019, 2021). Cultural traits play a role in the survival and reproduction of individuals and social groups; they reflect or even shape the structure of the population (Brakes et al., 2019), and can be lost when habitat fragmentation and population decline lead to reduced learning opportunities (Crates et al., 2021; Paxton et al., 2019). A comprehensive understanding of cultural change and loss, then, requires that we have the ability to detect and study not just intrinsic factorssocial, cultural, cognitivebut also extrinsic, ecological and demographic processes. This entails identifying the relevant spatial and temporal scales at which these processes manifest within natural populations, as well as their relative importance.

To contribute to this goal, we built a comprehensive dataset that spans three years and documents the dawn songs produced by male great tit birds during 454 breeding attempts in a single population located in Wytham Woods, UK. This population has marked variation in individual turnover, postnatal dispersal distances, age structure, and immigration across space (Figure 1), which allowed us to estimate their effects on song cultural repertoires at both individual and group levels within the long-term population study of this species (Lack, 1964). First, we assign more than 109,000 songs in 330 song repertoires to 242 individual birds through a combination of direct physical capture, radio frequency identification microchips, and a novel song-based reidentification method using a deep metric learning model.

**Figure 1.**
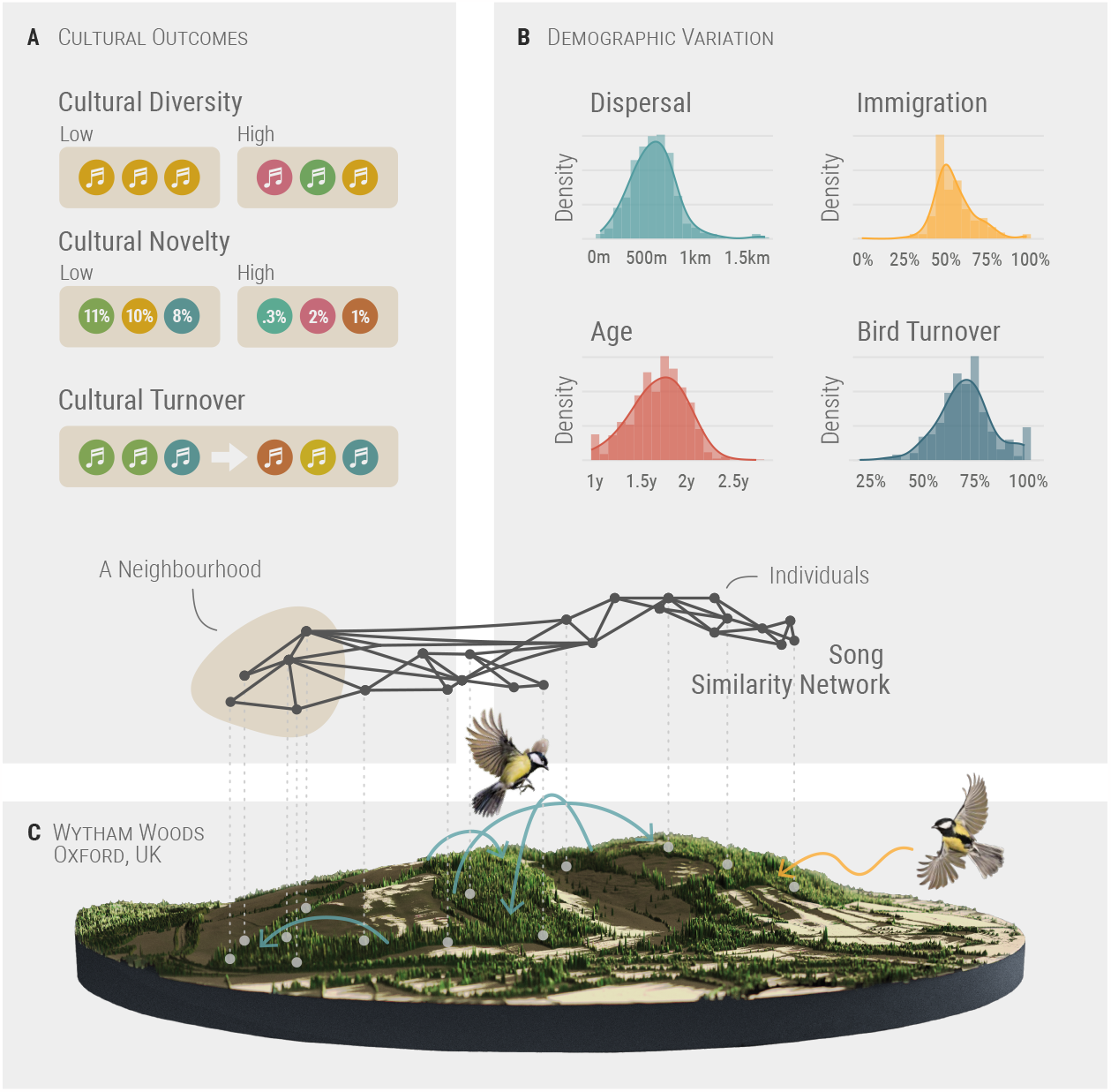
Study system and main variables in our analysis. (A) Cultural variables measured at the neighbourhood level. See methods for definitions. (B) Variation in the properties and composition of neighbourhoods across the population. See methods for definitions. (C) 3D render of our study site, Wytham Woods, based on first return LiDAR data (for Environment Food & Affairs, 2020) and made with rayshader (Morgan-Wall, 2023). Elevation is exaggerated. The network represents pairwise repertoire similarity between individuals with known spatial locations, used in the models reported in Fig. 2. Aquamarine (darker) arrows represent natal dispersal within the population, and the yellow (lighter) arrow represents immigration into the population, two of the variables used in this work. Age and bird (individual) turnover are not depicted.

Then we quantified individual and group-level traits and analysed variation in song cultural similarity, diversity, and turnover using network and spatially explicit Bayesian multilevel regression models.

Our results reveal an interplay of demographics and song cultural dynamics that, albeit complex, largely matches theoretical expectations, as discussed below. This work also demonstrates that bird song, which already provides what is perhaps the largest body of evidence for cultural change in animals (Laland & Janik, 2006), also has the potential to help us shed light on the impact of other population processes on animal cultures, owing to the fact that we can sample song cultural repertoires with relative ease.

## Reduced dispersal, increased immigration and an aged population are associated with higher cultural diversity

Population genetics provides robust evidence supporting the notion that high dispersal rates facilitate gene flow, which, in turn, reduces the efficacy of selection and diversification. Conversely, low dispersal facilitates genetic differentiation through mechanisms such as mutation and drift, leading to allopatric population divergence (Claramunt et al., 2011; Papadopoulou et al., 2009; Suárez et al., 2022). Drawing an analogy from genetics to culture, we anticipate that reduced dispersal rates will decelerate the diffusion of cultural traits (Nunn et al., 2009). This, in turn, should result in the maintenance of distinct behavioural patterns within populations (Planqué et al., 2014; Whitehead & Lusseau, 2012), leading to a greater abundance of cultural variants unique to a specific area. Our analysis indeed indicates that neighbourhoods where more birds have remained in proximity to their natal areas harbour greater cultural diversity and novelty (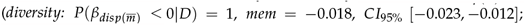;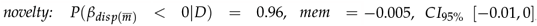 ; Figure 2A&B, Table S2), in line with prior research at a much coarser grain (Fayet et al., 2014).

**Figure 2.**
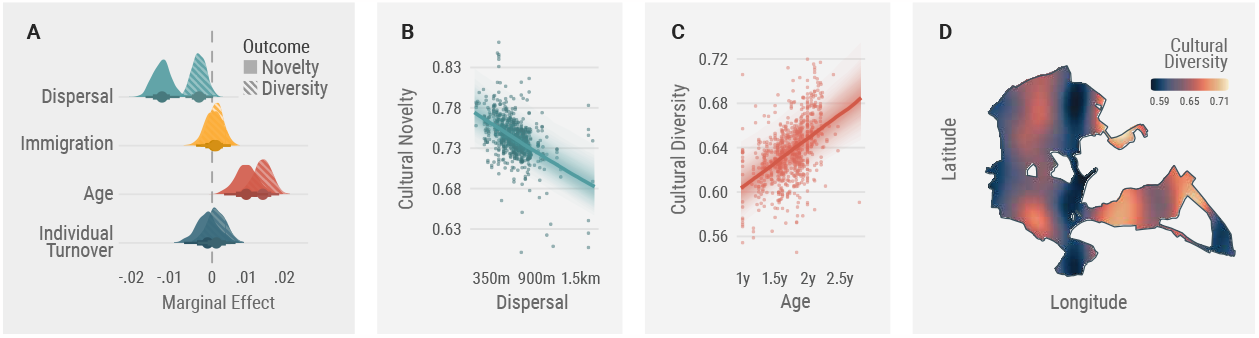
Influence of demographic variables on cultural diversity and novelty. (A) Marginal effects at the mean of neighbourhood characteristics including mean dispersal distance, proportion of immigrant birds, average age, and individual turnover. (B) Adjusted predictions and partial residuals of the effect of mean neighbourhood dispersal distance on cultural novelty. Low-dispersal neighbourhoods are those in which birds were born in the same area. (C) Adjusted predictions and partial residuals of the effect of the mean age of the neighbourhood on cultural diversity. A neighbourhood with a mean age of 1 would be one where all birds are breeding for the first time. (D) The average distribution of cultural diversity in the population across space during the study period (2020-2022). This map captures the residual variation in cultural diversity after taking into account the demographic variables in (A). The map is based on a Gaussian process model with an exponentiated-quadratic kernel covariance function, which allows us to interpolate between the locations where we have data.

However, the analogy breaks down when we examine our individual-level models more closely. Although the outcome at the group level resembles the homogenization of populations resulting from gene flow, the underlying mechanisms differ significantly, due to complex species-specific interactions between the timing of dispersal and learning mechanisms. In the case of great tits, these mechanisms are believed to involve selective retention or modification of songs encountered during early life and the establishment of territories following dispersal, a process that results in crystallised song repertoires that resemble those of their new neighbours at breeding sites (Marler & Peters, 1982; Nelson, 1992; Peters & Nowicki, 2017). Birds that dispersed over longer distances tend to have repertoires composed of songs that are common within the population (*novelty: P*(*β*_*disp*(*m*)_ *<* 0|*D*) = 1, *mem* = −0.2, *CI*_95%_ [−0.3, −0.09]; Figure 3B, Table S2), and possibly smaller repertoires as well (*rep*.*size: P*(*β*_*disp*(*m*)_ *<* 0|*D*) = 0.91, *mem* =−0.2, *CI*_95%_ [−0.44, 0.05]; Figure 3B; Table S2). We speculate that birds with more extensive movements are more likely to sample a larger proportion of common cultural variants, simply because they are exposed to more songs while dispersing. In contrast, birds with a more restricted and stable neighbour pool tend to be equally exposed to common and globally rarer songs, and this is sufficient to give rise to the differences that we detect at the group level (see repository for a simulation demonstrating this).

**Figure 3.**
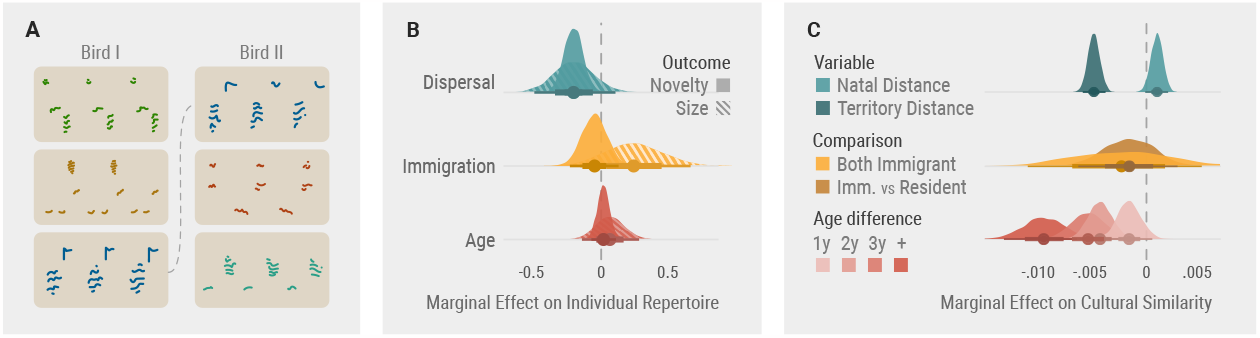
Individual and dyadic analysis of cultural diversity and similarity. (A) Illustrative example showing the repertoires of two different birds in the population, with three songs in each, one of which is shared. Each sub-panel shows a stylised spectrogram, with time in the horizontal axis and frequency in the vertical. Units not shown. (B) Marginal effects of dispersal, immigration status, and age on individual song repertoires, described in therms of their absolute size (the total number of distinct song variants sang by that bird) and their relative novelty (how frequent, on average, those songs are in the entire population within that year). (C) Marginal effects of dispersal, immigration status, and age on song repertoire similarity between individuals. Dispersal: Birds that are close neighbours are more culturally similar, regardless of where they were born, whereas natal distance may have a weak positive effect on cultural similarity. Immigration: There is no strong evidence that birds born outside the population are dissimilar from resident birds. Age difference: Birds are less culturally similar the greater the difference in their birth years.

Building on our understanding of cultural dynamics in relation to dispersal we expect that, when song learning is relatively precise and dispersal is limited, cultural differences will accumulate, and immigration will introduce cultural novelty to the recipient population. However, the extent to which immigration introduces new cultural variants into the population also hinges on an interplay between the species’ learning programme, the timing of dispersal, and the spatial movements of individuals. Animals that learn first and then disperse, for example, may bring cultural novelty with them. But this is not the case for great tits, whose young disperse in late summer and autumn, shortly after achieving independence; learn their songs until the end of their first winter (Rivera-Gutierrez et al., 2011), and become chiefly sedentary as adults (Dhondt, 1979; Dingemanse et al., 2003; Greenwood et al., 1979). In this species, then, we anticipate that immigrant birds will learn or retain songs they encounter upon arrival, either before or during the establishment of their territories (Graham et al., 2018; Keen, 2020).

Indeed, in our population, we find no evidence that the repertoires of birds originating from outside the population significantly differ from those of resident birds (*mem* = −0.002, *CI*_95%_ [−0.006, 0.002]; Figure 3C). This, in conjunction with the observation that cultural similarity between individuals is predicted by the distance between breeding territories (*mem* = −0.005, *CI*_95%_ [−0.006, −0.004]; Figure 3C; Table S2), supports the hypothesis that great tits are predominantly closed-end learners that learn primarily from territorial neighbours after dispersal (Graham et al., 2017; McGregor & Krebs, 1982b; Rivera-Gutierrez et al., 2011).

This leads to a somewhat contradictory scenario, however: immigrant birds, while not acoustically distinct, tend to exhibit larger repertoires compared to their resident counterparts (Figure 3B; 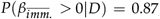, *mem* = 0.24, *CI*_95%_ [−0.098, 0.593]; Table S2). At the group level, this small and uncertain effect amplifies, such that neighbourhoods with a higher proportion of immigrant birds do not exhibit increased cultural diversity relative to the total number of songs (*mem* = 0.002 *CI*_95%_ [−0.004, 0.007]; Figure 2A); but they do have a higher absolute cultural diversity—above what would be expected based solely on the number of birds 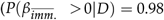, *mem* = 0.47, *CI*_95%_ [0.1, 0.84]; Figure S5, Table S2).

Previous research (Verhulst et al., 1997) has revealed that most birds arriving from outside the population disperse over two kilometres, significantly farther than the typical distances observed within the population (median for males = 558 metres Greenwood et al., 1979). This extended dispersal may have qualitative consequences for cultural diversity, through a combination of factors: first, an initial exposure to songs from the source population; then, a heightened pressure to adopt vocalisations similar to those of territorial neighbours to avoid any social or reproductive costs associated with non-local signals (Baker et al., 1981; Beecher, 2008; Lachlan et al., 2014; Mortega et al., 2014; Payne, 1983).

Finally, we find that individual turnover does not significantly affect cultural diversity or novelty, and we uncover an association between age structure and cultural diversity and novelty (Figure 2B). Individuals of the same generation share the most similar song repertoires and, while age itself doesn’t directly relate to changes in the repertoires of individual birds (Figure 3B), the acoustic similarity between pairs of individuals decreases as the age gap between them widens (Figure 3C; Table S2). This is expected when a species ceases to learn new songs as they age, and has detectable consequences for neighbourhoods: those with a higher proportion of older individuals have heightened levels of cultural diversity and novelty. Conversely, in areas where the majority of the population comprises active learners surrounded by their peers, birds tend to produce fewer unique songs that are simultaneously more common within the population (Figure 2A; Figure S5; 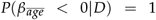, *mem* = 0.021, *CI*_95%_ [0.014, 0.027]; 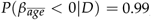 0.99, *mem* = 0.012, *CI*_95%_ [0.005, 0.019]).

### Demographic processes strongly moderate the rate of cultural change at small spatiotemporal scales

We now shift our focus from static measures of cultural diversity to cultural turnover, examining how quickly song variants disappear from neighbourhoods and the consequences this has for their cultural makeup. The primary driver of cultural turnover is individual turnover (total effect *mem* = 0.072 *CI*_95%_ [0.051, 0.093]): as birds leave or die, many song variants disappear with them. Accounting for this, we also assess the direct impact of mean natal dispersal distance, the proportion of immigrant birds, and mean neighbourhood age: higher levels of each of these factors correlate with slower cultural change in the neighbourhood (Figure 4A; Table S2). When there is substantial dispersal, a high influx of immigrants, and an age distribution skewed towards older individuals, the model predicts slower cultural change, at less than half the rate compared to the converse scenario (0.28 *CI*_95%_ [0.23, 0.34] vs. 0.61 *CI*_95%_ [0.49, 0.76], as illustrated in Figure 4E). Modelling work suggests that learning from older individuals should slow down cultural change (Kirby & Tamariz, 2021), aligning with our observations (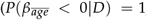. Age may serve as a brake on change, potentially increasing the relative cultural diversity and novelty within neighbourhoods by maintaining song types now less frequent in the population, as supported by the individual-level analysis where birds become more dissimilar as they are further in time. Across the three-year study period, now considering the entire population, cultural turnover between consecutive years hovers around 45% (0.47, 0.44). If all variants faced an equal chance of disappearing, this high turnover rate would lead to complete cultural replacement within a short time span. However, with a two-year gap, turnover only slightly increases to 0.59. We expect this rate to taper over longer periods, as rare variants encounter greater stochasticity while common songs endure (Figure S4A). This is exemplified by some common song types documented over four decades ago that persist within the same population (Keen, 2020; McGregor & Krebs, 1982b), either through accurate learning or, more likely, strong convergent biases (Claidière & Sperber, 2007; James & Sakata, 2017; Lachlan et al., 2018; Tchernichovski et al., 2021).

**Figure 4.**
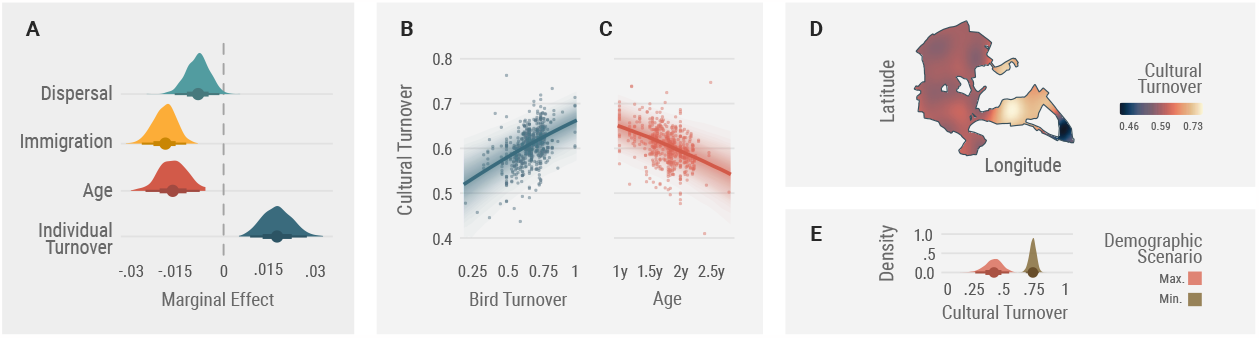
Influence of demographic variables on the rate of local cultural change. (A) Marginal effect at the means for mean dispersal distance, proportion of immigrant birds, average age, and individual turnover on the rate of cultural turnover. (B and C) Adjusted predictions and partial residuals of the effects of the proportion of individual turnover on cultural turnover (B) and the effect of mean neighbourhood age on cultural turnover (C). (D) The population’s average distribution of cultural turnover across space during the study period (2020-2022). (E) Posterior counterfactual predictions for two scenarios: all variables at their maximum (Max.) and minimum (Min.) observed values, adjusting for individual turnover. Cultural turnover is expected to be over two times higher when neighbourhood dispersal, immigration and age are low.

### Consequences for cultural structure, stability and diversity

Cultural traits, learnt bird song in this case, are shaped by many factors: some external, such as those discussed here, others intrinsic to learning and culture, and yet others that arise from selective processes driven by preference and function. Even within the confines of a relatively small population— Wytham Wood spans a mere four kilometres—we have recovered associations between heterogeneity in the demographic composition of neighbourhoods and cultural outcomes. This emphasises the need for both empirical studies and modelling efforts on cultural change to account for the population’s demographic characteristics and their inherent heterogeneity across time and space, which shape individuals’ exposure to cultural variants and opportunities for learning and, therefore, emergent group-level cultural dynamics.

## METHODS

### Resource availability

The complete Wytham great tit song dataset is available in osf.io/n8ac9, and documented here. The main repository with code and data to reproduce the analysis and figures in this article can be found at birdsong-demography.

### Data collection

#### Study system and fieldwork

Great tits are small, short-lived birds—average lifespan: 1.9 years—that sing acoustically simple yet highly diverse songs. Each male great tit has a repertoire of one to over 10 song variants, referred to as ‘song types,’ which are repeated multiple times in short bursts separated by longer periods of silence. During the breeding season, from March to June, great tit pairs are socially monogamous and defend territories around their nests (Hinde, 1952). In Wytham Woods, Oxfordshire, UK (51*°*46 N, 1*°*20 W), a population of these birds has been the focus of a long-term study since 1947 (Lack, 1964). Wytham Woods is a semi-natural predominantly deciduous woodland that spans an area of approximately 385 hectares and is surrounded by farmland. Most great tits in this population breed in nest boxes with known locations, and the majority of individuals are marked with a unique British Trust for Ornithology (BTO) metal leg ring as either nestlings or adults.

We collected data from late March to mid-May during the 2020, 2021, and 2022 breeding seasons. Every year, fieldworkers checked each of the 1018 nest boxes at least once a week before and during the egg-laying period, which typically lasts from one to 14 days (Perrins, 1965), and recorded the identities of breeding males and females, the dates of clutch initiation and egg hatching, clutch size, and fledgling number and condition under standardised protocols. We found the first egg date by assuming that one egg is laid every day and counting back from the day of observation. In cases where we did not observe the chicks on their day of hatching, the actual hatching date was determined by assessing the weight of the heaviest chicks and extrapolating their age from established growth curves (Cresswell & Mccleery, 2003; Gibb, 1950).

To record the vocalisations of male great tits we took advantage of their behaviour during the reproductive period, when they engage in continuous singing near their nests at dawn before and during egg laying (Mace, 1987). Collectively, this vocal display is referred to as the dawn chorus, and has been demonstrated to yield a reliable estimation of the song repertoire of individuals when recorded in full (Rivera-Gutierrez et al., 2012; Van Duyse et al., 2005). As soon as we suspected that a pair of great tits were using a nest box—based on nest lining materials, egg size if present, or other signs of activity— we deployed an autonomous sound recorder nearby. Our goal was to maintain a consistent position and orientation for the recorder. The microphone faced upwards and slightly away from the nest box, aligning with the nest box entrance hole’s direction. The birds sang near the recorder, and although we did not gather data on this aspect, our anecdotal observations were in line with a different population where the average distance to the nest box was 10 metres (Halfwerk et al., 2012). The birds also changed perches and moved around during our recording. Although variation in sound amplitude due to changes in distance and direction could affect song selection, we did not observe systematic bias, ruling out potential issues like consistently low signal-to-noise ratios causing exclusion of entire song types.

All work involving birds was subject to review by the University of Oxford, Department of Zoology, Animal Welfare and Ethical Review Board (approval number: APA/1/5/ZOO/NASPA/ Sheldon/TitBreedingEcology). Data collection adhered to local guidelines for the use of animals in research and all birds were caught, tagged, and ringed by BTO licence holders (NMR’s licence: C/6904).

#### Recording equipment and schedule

We used 60 (30 in 2020) AudioMoth recorders (Hill et al., 2019), which were housed in custom-built waterproof enclosures. Recording began approximately one hour before sunrise (05:36 – 04:00 UTC during the recording period) and consisted of seven consecutive 60-minute-long recordings with a sample rate of 48 kHz and a depth of 16-bit. To sample as many birds as possible, we left each recorder at the same location for at least three consecutive days before moving it to a different nest box. We relocated 20 recorders (10 in 2020) every day throughout the recording period.

#### Data processing and annotation

We processed and annotated the song recordings, 109,963 in total, using custom software and scripts written in Python 3 (van Rossum, 1995) and the open source package pykanto (Merino Recalde, 2023a). These are available from github.com/nilomr/greattit-hits-setup (Merino Recalde, 2023b). Our annotated dataset and a detailed description of the process can be found in Merino Recalde et al. (2023).

#### Identifying individuals and their traits

We further augmented our dataset by training a deep metric learning model (see (Merino Recalde et al., 2023) for details) to recognize individual songs, which we then used to assign individual IDs to a subset of birds that we failed to physically capture or identify using PIT (Passive Integrated Transponder) tags. This increased the number of identified breeding attempts for which we also had songs from 299 to 330, belonging to 242 unique birds. Briefly, we calculated pairwise song distances using the feature vectors obtained from the trained model. Then we assigned unknown song repertoires to known birds if they met two conservative criteria: that at least two songs had a Euclidean distance below 0.9, and that the unknown singer was recorded less than 100 metres apart from the known individual (see Figure S3 for a graphic explanation). Natal dispersal distance was calculated as the straight line distance from the natal site to the breeding site. The dispersal distances of birds classed as immigrants (not ringed as chicks in the population) are not known, but most are thought to come from other populations at least 1 km, and likely more than 2.5 km, away (Quinn et al., 2011; Verhulst et al., 1997). We determined age based on the year of hatching for birds born in the population; and plumage characteristics for immigrants, which are most often caught as yearlings (76%)—allowing us to age them accurately (Woodman et al., 2023).

#### Characterising repertoire similarity

Our analyses require i) a measure of the acoustic similarity between any two birds, and ii) a way to identify song cultural variants. The underlying assumption is that song repertoires will be more similar if one bird has learned it at least in part from a second, or if they have both learnt from other individuals who are themselves similar due to intergenerational cultural descent. There is no single optimal solution for this problem, both due to technical challenges and because we do not know enough about song perception and learning mechanisms in this species. There are three main possible approaches, each with its own advantages and disadvantages.

#### Continuous similarity

Traditional methods used to compare bird vocalisations include visual inspection of spectrograms and measurement of hand-picked acoustic features. However, these approaches have limitations in dealing with noise and variations in performance and can be extremely time-consuming. They also fail to capture complex features such as the syntactic relationships between notes. So, instead, we adopted a data-driven approach by training a Vision Transformer (ViT) model for feature extraction in a metric learning task. Our goal was to create a similarity space based on inherent variation in the data, using categorical labels of song types sung by individual birds, which we know to be perceptually and behaviourally significant (Lind et al., 1996). Further details and code are available at (Merino Recalde, 2023a) and (Merino Recalde et al., 2023). We used the resulting model to calculate feature vectors for each song in the dataset (109,963 samples x 384 dimensions), which serve as compressed representations that can be used to compare them.

Great tits have variable repertoire sizes and there is no evidence that they ever learn them en bloc (McGregor & Krebs, 1982b; Rivera-Gutierrez et al., 2010). Therefore, the simplest continuous measure (an average pairwise Euclidean distance between all songs) would mask any signatures of learning if the average repertoire similarity is similar across the population, and does not take into account the asymmetry in total repertoire size. To improve on this, we define repertoire similarity as the average minimum Euclidean distance (AMED), given by

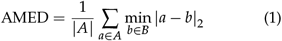

where we compare each song feature vector *a* in set *A* with all song feature vectors *b* in set *B* and compute their Euclidean distance |*a* − *b*|_2_. We then retain the minimum distance for each element in set *A* and obtain the AMED by averaging these minimum distances over all elements in set *A*. The main advantage of this approach is that it allows us to avoid imposing discrete population-wide song categories. On the other hand, if song learning is categorical and not very precise in terms of fine song structure, this method could underestimate it or fail to detect it. We used this approach for all individual-level analyses in this paper.

#### Automated clustering

Instead of calculating a continuous measure of repertoire similarity, we can build a pairwise distance matrix for all songs, assign them to discrete clusters using a clustering algorithm, and then calculate the intersection between repertoires by using the Jaccard coefficient or modelling it as a binomial process, with *n* = the combined repertoire size and *s* = the number of songs in the same cluster. Here we used UPGMA hierarchical clustering and dynamic tree-cut techniques to classify the syllables into distinct types, allowing a minimum cluster size of 1 to ensure the representation of rare song types. The usefulness of this method relies on the global properties of the embedding space derived from section **??**. In a low-dimensional space where linear distances effectively capture meaningful variation, creating clusters by cutting the hierarchical tree at different heights yields varying cluster counts while maintaining meaningful groupings. However, in a high-dimensional space where global distances are not meaningful, only relatively small clusters of nearby points remain interpretable. This is the case with our dataset and embedding space: we find that the method reliably groups song renditions by the same bird across different years, alone or together with other birds with highly similar songs, yet consistently splits songs that are similar by human (and perhaps great tit (Falls et al., 1982)) standards, ultimately leading to a very large number of clusters (the most stable clustering solutions were close to the total number of different individual song types, >1000). Due to these issues, we did not use song types defined in this way.

#### Manual categorization

To date, all research on great tit song has relied on a visual classification of songs into population-level types (Baker et al., 1987; Falls et al., 1982; Fayet et al., 2014; Hutfluss et al., 2022; McGregor & Krebs, 1982a; McGregor & Krebs, 1982b; McGregor et al., 1981). This process is both inevitable and very subjective. However, despite its clear problems, human perceptual judgments might be our best available substitute for those of the birds (but see recent work by Morfi et al., 2021; Zandberg et al., 2022) for some tasks. Indeed, across fields, advanced classification algorithms are often evaluated against ground truth created by humans, and this is also the case in bird song research.

Our neighbourhood-level analyses require that we define discrete cultural units, so, given the difficulties with the alternatives described above, we adopted a variant of this approach and used the criteria followed by McGregor and Krebs (1982b) and most subsequent work. With over 100,000 songs, our dataset is much larger than is common in the field and would have been impossible to label entirely manually. Instead, we used the output of the process described above, consisting of labelled song repertoires (birdID x song type). This made the problem 57 times smaller: 1920 song variants that were already assigned to small clusters of highly similar songs, which we reviewed manually.

Following common practice in the field, we validated our manually assigned labels statistically, although we note that i) the ability of a statistical method to differentiate between manually defined clusters does not mean that these are perceptually meaningful, only that they can be distinguished in a manner that aligns with human classification, and ii) a large range of clustering solutions will be compatible with the data. To do this, we retrained the ResNet50-based classifier described in Merino Recalde (2023a) using a random subset of the data and obtained an accuracy of 0.87 on the validation set (see other metrics in the repository). With the caveats already mentioned, this means that our manual classification following McGregor and Krebs (1982b) is successful at finding a stable solution that reduces intraclass variation. A comparable process by Fayet et al. (2014) was able to reach 0.71 accuracy for 374 songs. We further explored the result by building a dendrogram based on the confusion matrix during test time and reviewing the classes that were not well supported, which led us to combine seven classes into two. There is an inverse relationship between how densely occupied a region of the song space is and the ease with which we can find categorical divisions: the more examples the more graded the variation and, in consequence, what may have seemed like clear-cut categories if we had fewer data blend into one another without an obvious transition.

In practical terms, because most of the great tits in our population sing some variation of the well-known ‘tea-cher, tea-cher’ song, these are much harder to categorize than the many rare songs with complex structures only sung by one or a few birds. This was our impression when manually labelling the songs, and it was also the case when applying the supervised classification algorithm. As mentioned in the main text, the direct consequence of this for our analysis is that the absolute estimates of cultural turnover depend on the granularity of this process: when we lump all similar ‘tea-cher’ songs, as McGregor and Krebs (1982b) do, the estimates of turnover are necessarily lower—but, crucially, any relative differences remain the same. The code used to perform the song type validation process, along with the figures generated during it, can be found in the main narrative notebook and a dedicated repository.

### Quantification and statistical analysis

#### Pairwise similarity and individual repertoire models

It is common for analysis of song similarity to fit simple linear regression models using all pairwise comparisons in a population. However, this leads to very strong pseudo-replication and, therefore, an increased chance of Type I errors. To avoid this, we treat our song similarity data as a fully connected network and build Bayesian multilevel models with a multi-membership structure and the pairwise AMED described above as the response variable. The full model specifications can be found in the main repository for this project; also see a summary in Table S1.

#### Individual repertoires

We first modelled individual repertoire size using Poisson and negative binomial models, but this led to poor performance as assessed through posterior predictive checks (both underestimation of mean values and either under or over-estimation of very low repertoire sizes). Instead, we built continuation ratio models, a type of sequential ordinal model where reaching a particular level (number of song types in the repertoire) requires first reaching all lower levels (Chambers & Drovandi, 2023; Warti et al., 2020). 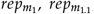 and 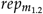 estimate the association between immigrant status, distance dispersed, age, and repertoire size. Three further log-normal models, 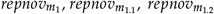, do the same for the average cultural diversity of individual repertoires

#### Pairwise similarity

Our first model 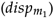 explores the interaction between natal distance, that is, the distance between the nests where two resident birds were born, and the distance between the centre of their breeding territories, adjusting for year and absolute age difference. We do not have direct information on how long birds have spent around one another, so instead we estimate the effect of the interaction of the distance at which they were born and the distance at which they subsequently breed: If both are small, they will have had more opportunities for interaction and learning. We extract predictions for the interaction and calculate marginal effects at minimum distances, to answer the questions ‘How does cultural similarity change with distance for birds that were born nearby’ and ‘Does how close a bird was born matter for birds that hold territories nearby’. We use a similar model structure 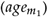 to estimate the marginal effects of the absolute age difference, this time adjusting for the natal and territorial distance between birds. Then, to study the effect of immigration, we fit a model 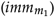with the possible combinations of immigration status (both immigrant, both residents, one of each) and adjust for age difference and territorial distance.

### Group-level properties

#### Defining neighbourhoods and their demographic properties

Song turnover, diversity, and novelty are group-level properties. However, our study lacks naturally occurring distinct subpopulations that we can use as units for analysis. Rather than partitioning the population using a discrete polygonal grid or non-overlapping areas, we opted to model neighbourhoods continuously across space, with a radius of 200 m around each nest box occupied at least once during the study (Fayet et al., 2014) which we sampled across the duration of the study. This radius is necessarily arbitrary but strikes a good compromise between describing the relevant spatial scale at which vocal interactions occur, which extends up to around 180 metres (Bircher et al., 2021; Blumenrath & Dabelsteen, 2004), and maintaining an adequate sample size in areas of low density. Neighbourhoods defined in this way are highly non-independent, so we model both this methodological spatial dependence and other sources of spatial autocorrelation intrinsic to the study site by including a 2D Gaussian process (GP), which estimates a length-scale parameter defining a variance-covariance matrix for the spatial locations based on their distance (Dearmon & Smith, 2016; Gelfand & Schliep, 2016; Wright et al., 2021). We confirmed that this eliminated the residual spatial autocorrelation via Moran’s I tests. Note that we fit a separate GP for each year, as treating the spatial dependence as fixed across the study duration, as is often done, risks further underestimating uncertainty.

We define our predictor variables in the following way: Individual turnover is the proportion of birds that were not already in a neighbourhood in the preceding year 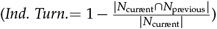. Dispersal is the mean of the distances that birds in the neighbourhood travelled to get from their natal territories to their breeding territories if they were born within the Wytham population. Immigration is the proportion of birds that were not ringed as nestlings in the population, and neighbourhood age is the mean age of the birds within it. Figure S1 illustrates that our sampling process did not introduce bias into any of these predictor variables: the birds from which we recorded song repertoires were, on average, representative of the true neighbourhood composition.

#### Operational definitions of cultural diversity, novelty, and turnover

We calculated a simple diversity index by dividing the number of different song types by the total number of songs in a neighbourhood. To calculate the novelty index, we computed the relative frequency of each class label in the current year in the entire population. We then took the mean of these relative frequencies for each song type in the neighbourhood, took the logarithm of the inverse of this proportion and scaled it between 0 and 1. In this way, ‘diversity’ describes the proportion of unique songs in a neighbourhood, and ‘novelty’ refers to how uncommon, on average, the songs of the birds in a neighbourhood are. These two ways of characterising cultural diversity are (as expected) anti-correlated in our study site due to the effect of sampling: more frequent songs are sampled more readily, causing larger sample sizes (neighbourhoods with more density and therefore songs) to yield lower average estimates of diversity and higher average estimates of novelty, in a nonlinear manner. Once this is adjusted for, diversity and novelty are positively correlated, as expected (see Figure S2; models 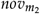 and 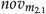). All of our models adjust for this sampling effect.

#### Models

To study the effect of dispersal and immigration on local cultural diversity and novelty, we built log-normal models 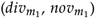 and estimated the marginal effects of the proportion of immigrants, mean dispersal distance, and mean neighbourhood age, while also adjusting for individual turnover, year, and spatial dependence. Lastly, to examine whether the effects of immigration and dispersal on cultural diversity were related to individual differences in repertoire size and novelty, we fit two further models predicting the absolute number of unique songs in a neighbourhood while also adjusting for the number of birds 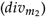 and the number of songs 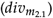.

We calculated the rate of song cultural turnover as the proportion of unique song types in a given year that were not already present in the same neighbourhood the preceding year, and this was the response variable in two models: one 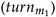 trying to estimate the total effect of turnover and a second 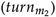 estimating the marginal effects of the proportion of immigrants, mean dispersal distance, and mean age while also adjusting for individual turnover, year, and spatial dependence. In both cases, we modelled the response distribution as a truncated log-normal with a hurdle (logistic) part to account for the zeroes.

##### Model estimates and reporting

We build the models and approximate the posterior distributions of the parameters of interest using brms (Bürkner, 2017), an interface to the Hamiltonian Monte Carlo engine Stan (Stan Development Team, 2023). We then processed the posterior distributions with the help of the marginal effects package. We checked model convergence via the effective number of samples, visual inspection of the chain trace plots, and the Gelman-Rubin diagnostic. Estimation in a Bayesian framework returns a posterior distribution of possible values instead of point estimates. By convention, we report posterior central estimates (means or medians) and their 95% credible intervals, but also include plots with full posteriors. Note that categorical predictors are dummy-coded and continuous predictions z-score transformed.

For each parameter of interest, we calculate predictions or marginal effects at the means or other relevant values. Regression plots show predicted values of the mean and their credible intervals, as well as partial residuals adjusted to the means or other relevant values of the explanatory terms included in the model (Fox & Weisberg, 2018; Larsen & McCleary, 1972). We have tried to build reasonable models, but even then our estimates should not be interpreted causally. See the software section at the end for a complete list of libraries used in the various analyses and the code repository for full model specifications.

## ACKNOWLEDGEMENTS

We thank all those who have contributed to the long-term nest box study in Wytham Woods and the collection of associated data. This work was supported by a Clarendon-Mary Frances Wagley Graduate Scholarship and an EGI scholarship to Nilo Merino Recalde, and made use of the University of Oxford Advanced Research Computing facility (Richards, 2015).

## AUTHOR CONTRIBUTIONS

**Nilo Merino Recalde**: Conceptualization, Methodology, Software, Formal Analysis, Investigation, Data Curation, Writing - Original Draft, Writing - Review & Editing, Visualization. **Andrea Estandía**: Investigation, Data Curation, Writing - Review & Editing. **Sara C. Keen**: Writing – Review & Editing. **Ella F. Cole**: Supervision, Project Administration. **Ben C. Sheldon**: Supervision, Project Administration, Writing – Review & Editing, Funding Acquisition.

## SUPPLEMENTARY INFORMATION

**Table S1.**
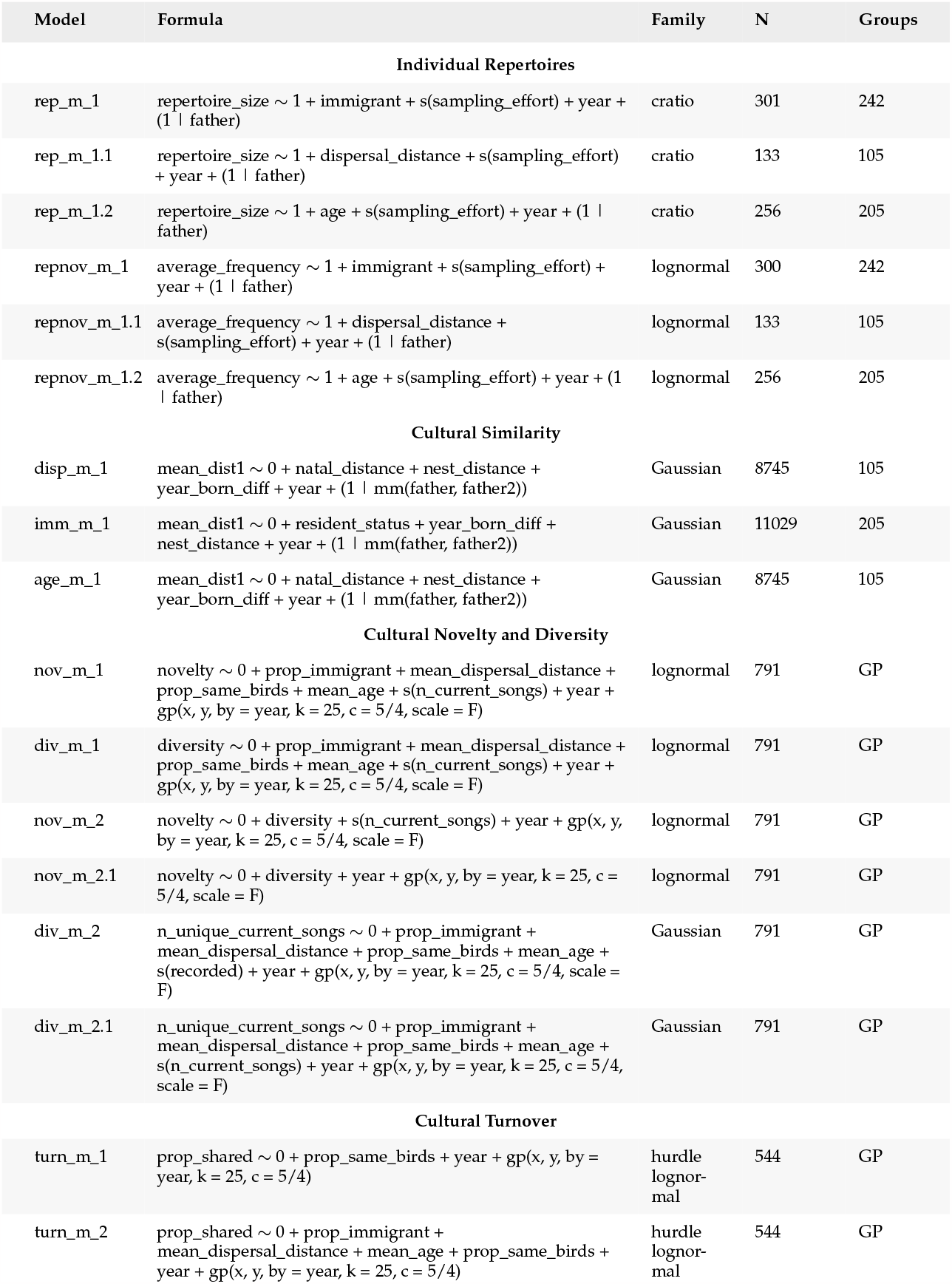
Model information

**Table S2.** Model estimates

| Model | Hypothesis | Estimate<sup>a</sup> | Evid. Ratio | Post. Prob |
| --- | --- | --- | --- | --- |
| **Individual Repertoires** |  |  |  |  |
| rep_m_1 | immigrant > 0 | 0.239 [-0.098, 0.593] | 6.963 | 0.874 |
| rep_m_1.1 | dispersal distance < 0 | -0.201 [-0.443, 0.045] | 10.111 | 0.910 |
| rep_m_1.2 | age > 0 | 0.064 [-0.108, 0.241] | 2.701 | 0.730 |
| repnov_m_1 | non-immigrant < 0 | -0.049 [-0.2, 0.1] | 2.401 | 0.706 |
| repnov_m_1.1 | dispersal distance > 0 | 0.203 [0.088, 0.316] | 741.857 | 0.999 |
| repnov_m_1.2 | age > 0 | -0.017 [-0.093, 0.058] | 0.540 | 0.351 |
| **Cultural Similarity** |  |  |  |  |
| disp_m_1 | natal distance > 0 | 0.001 [0, 0.002] | 22.529 | 0.958 |
| disp_m_1 | nest distance > 0 | -0.005 [-0.006, -0.004] | Inf | 1 |
| imm_m_1 | both resident-both immigrant < 0 | 0.002 [-0.005, 0.01] | 0.438 | 0.304 |
| imm_m_1 | both resident-one immigrant < 0 | 0.002 [-0.002, 0.006] | 0.331 | 0.248 |
| age_m_1 | age difference 0-1 > 0 | 0.002 [0, 0.003] | 12.289 | 0.925 |
| age_m_1 | age difference 0-2 > 0 | 0.004 [0.002, 0.006] | Inf | 1 |
| age_m_1 | age difference 0-3 > 0 | 0.006 [0.003, 0.008] | 1999 | 1 |
| age_m_1 | age difference 0-4+ > 0 | 0.01 [0.006, 0.013] | Inf | 1 |
| **Cultural Novelty and Diversity** |  |  |  |  |
| nov_m_1 | mean dispersal distance < 0 | -0.018 [-0.023, -0.012] | Inf | 1 |
| nov_m_1 | proportion immigrant < 0 | 0.001 [-0.005, 0.006] | 0.752 | 0.429 |
| nov_m_1 | mean age < 0 | 0.012 [0.005, 0.019] | 399 | 0.998 |
| nov_m_1 | individual turnover < 0 | 0.001 [-0.005, 0.008] | 1.733 | 0.634 |
| div_m_1 | mean dispersal distance < 0 | -0.005 [-0.01, 0] | 26.65 | 0.964 |
| div_m_1 | proportion immigrant < 0 | 0.002 [-0.004, 0.007] | 0.442 | 0.306 |
| div_m_1 | mean age < 0 | 0.021 [0.014, 0.027] | Inf | 1 |
| div_m_1 | individual turnover < 0 | -0.002 [-0.008, 0.005] | 0.446 | 0.309 |
| nov_m_2 | diversity > 0 | 0.713 [0.629, 0.795] | Inf | 1 |
| nov_m_2.1 | diversity > 0 | -0.099 [-0.216, 0.018] | 0.086 | 0.080 |
| div_m_2 | mean dispersal distance < 0 | -0.658 [-0.999, -0.315] | 570.429 | 0.998 |
| div_m_2 | proportion immigrant < 0 | 0.469 [0.1, 0.837] | 62.492 | 0.984 |
| div_m_2 | mean age < 0 | 1.045 [0.576, 1.495] | Inf | 1 |
| div_m_2 | individual turnover < 0 | 0.204 [-0.291, 0.683] | 3.077 | 0.755 |
| div_m_2.1 | mean dispersal distance < 0 | 0.019 [-0.213, 0.249] | 0.824 | 0.452 |
| div_m_2.1 | proportion immigrant < 0 | 0.072 [-0.168, 0.312] | 2.259 | 0.693 |
| div_m_2.1 | mean age < 0 | 0.928 [0.599, 1.245] | Inf | 1 |
| div_m_2.1 | individual turnover < 0 | -0.031 [-0.349, 0.279] | 0.726 | 0.421 |
| **Cultural Turnover** |  |  |  |  |
| turn_m_1 | individual turnover > 0 | 0.072 [0.054, 0.09] | Inf | 1 |
| turn_m_2 | mean dispersal distance < 0 | -0.022 [-0.039, -0.006] | 79 | 0.988 |
| turn_m_2 | proportion immigrant < 0 | -0.051 [-0.065, -0.037] | Inf | 1 |
| turn_m_2 | mean age < 0 | -0.044 [-0.063, -0.026] | 3999 | 1 |
| turn_m_2 | individual turnover < 0 | 0.047 [0.028, 0.066] | Inf | 1 |

<sup>a</sup> Estimates are Medians and 95% Credible Intervals

**Figure S1.**
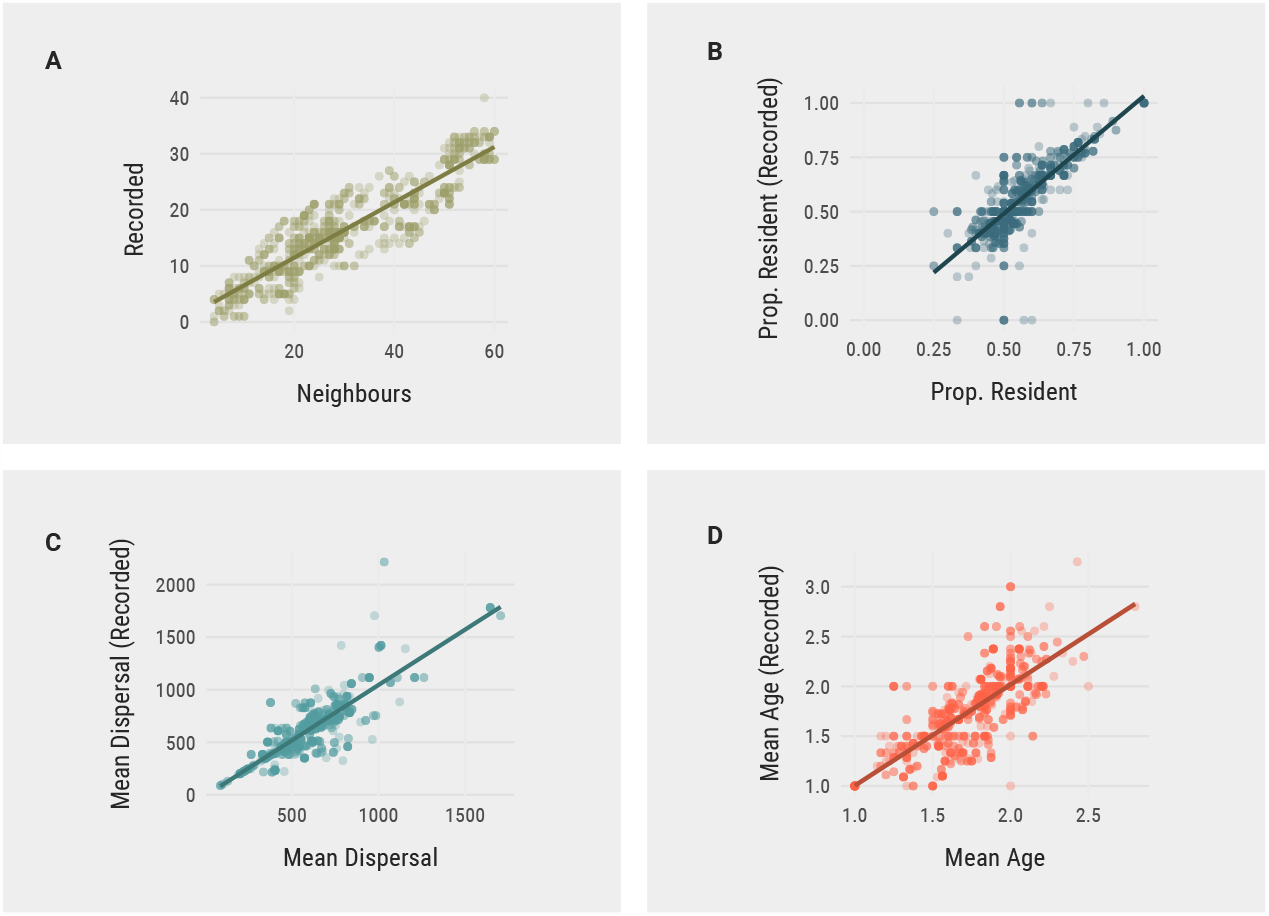
Absence of bias in the sampling of neighbourhood properties. Correlation between the actual neighbourhood properties and neighbourhood properties estimated from birds for which we have song recordings. (A) Neighbourhood size (number of individuals) and number of individuals with song recordings. (B) Proportion of resident birds from monitoring data and only those birds with song recordings. (C) Mean dispersal distance calculated from birds born in the study site and only those birds born in the study site with song recordings. (D) Mean age of birds in the study site and only those birds with song recordings.

**Figure S2.**
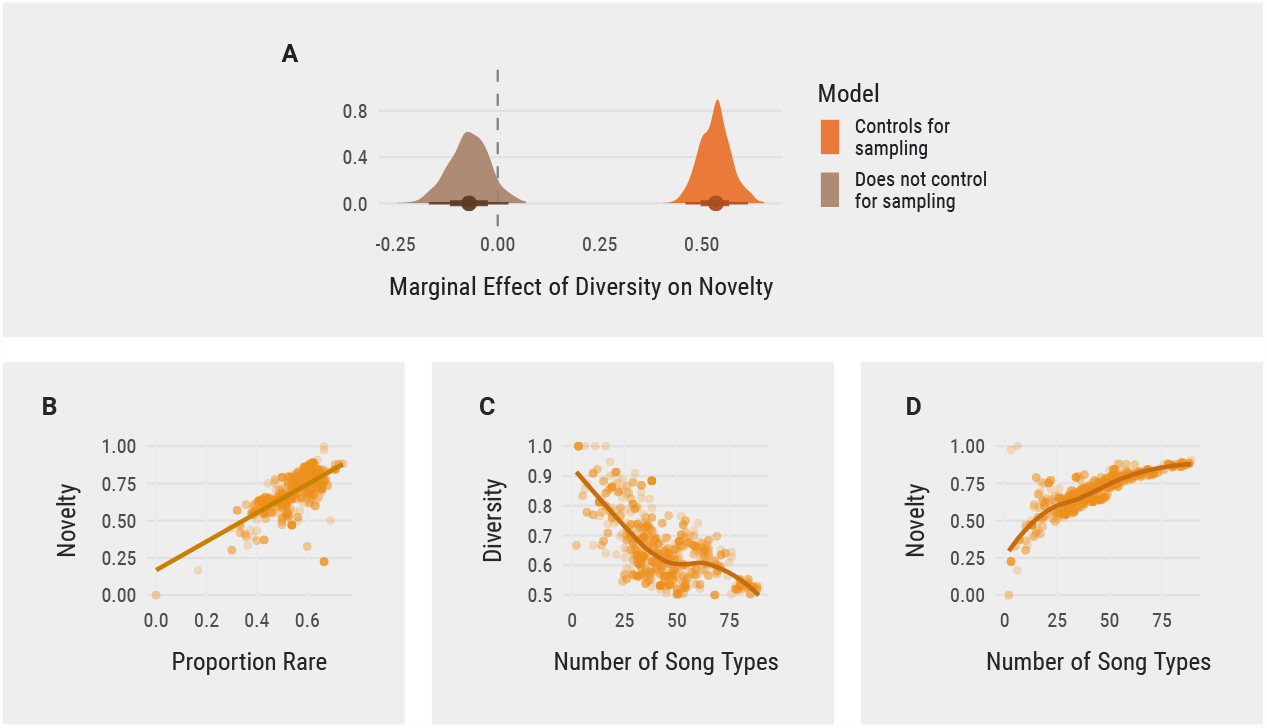
Relationships among outcome variables and sampling effects. (A) Marginal effect of diversity—which describes the proportion of unique songs in a neighbourhood—on novelty, that is, how uncommon, on average, the songs of the birds in a neighbourhood are. These two ways of characterising cultural diversity are (as expected) anti-correlated in our study site due to the effect of sampling: more frequent songs are sampled more readily, causing larger sample sizes (neighbourhoods with more birds and therefore songs) to yield lower average estimates of diversity (C) and higher average estimates of novelty (D), in a nonlinear manner. Once this is adjusted for, diversity and novelty are positively correlated, as expected. (B) Our measure of cultural novelty (y-axis) has the advantages of being continuous and not using an arbitrary cutoff, but is nonetheless correlated with definitions traditionally used in the literature, such as ‘songs shared by fewer than 4 birds (McGregor & Krebs, 1982b)

**Figure S3.**
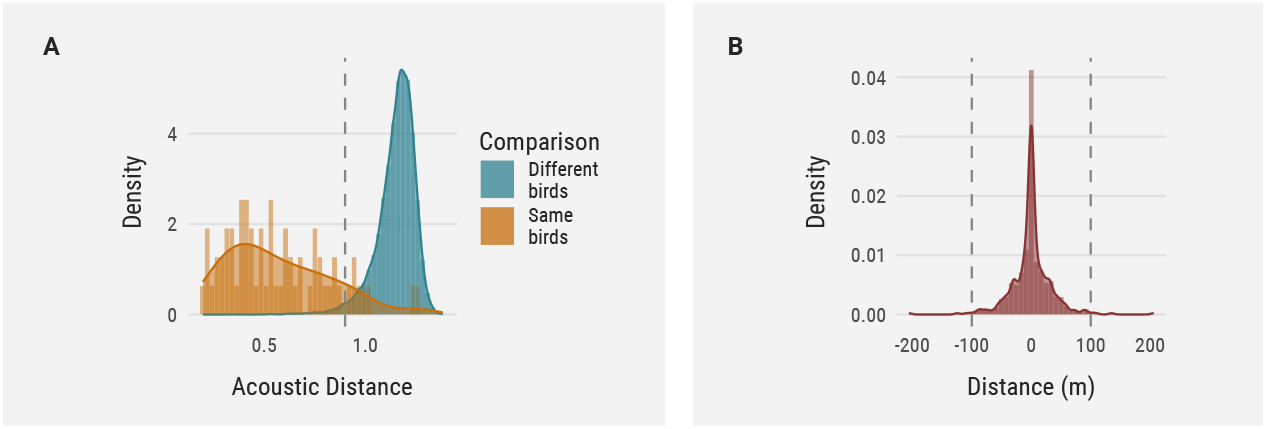
Thresholds used during the process of reidentifying individual birds based on their songs. (A) Shows the distribution of the acoustic distances between the same song type sung by the same known bird in different years, in orange, and the minimum pairwise distance between different birds and years. The x-intercept of the vertical line = 0.9. (B) Shows the distribution of the change in distance from the natal nest box to the breeding site in different years for birds that bred more than once. Adult birds have high nest site fidelity, which we used as a further constraint when reidentifying them from their songs.

**Figure S4.**
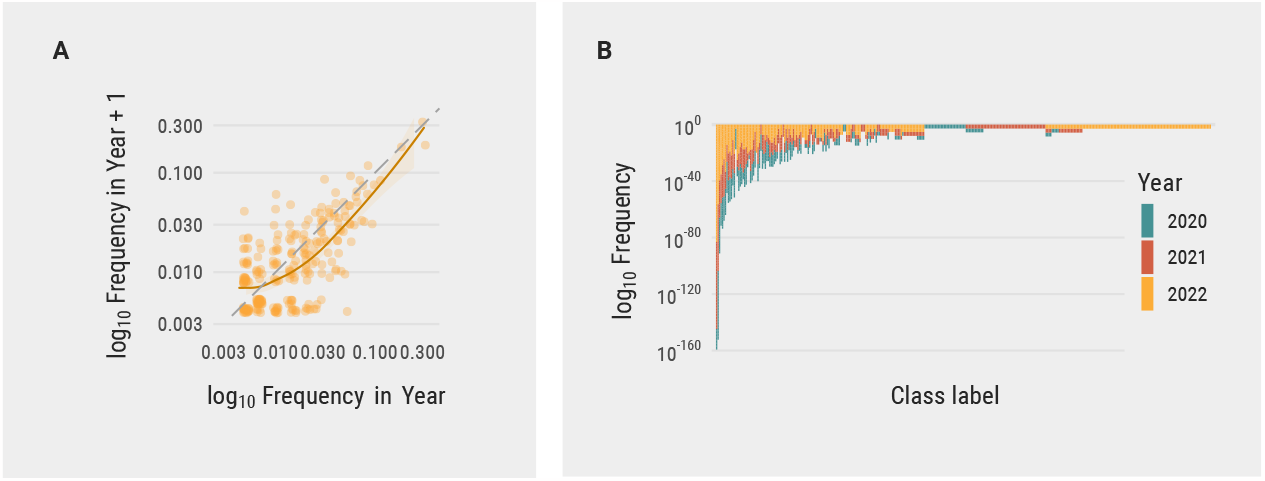
Song frequencies and their relationship with abundance in the following year. **(A)**The abundance of a song type in a given year predicts its abundance in the following year, with higher variance around rare songs. (B)Histogram showing the frequency of individual song types in the study.

**Figure S5.**
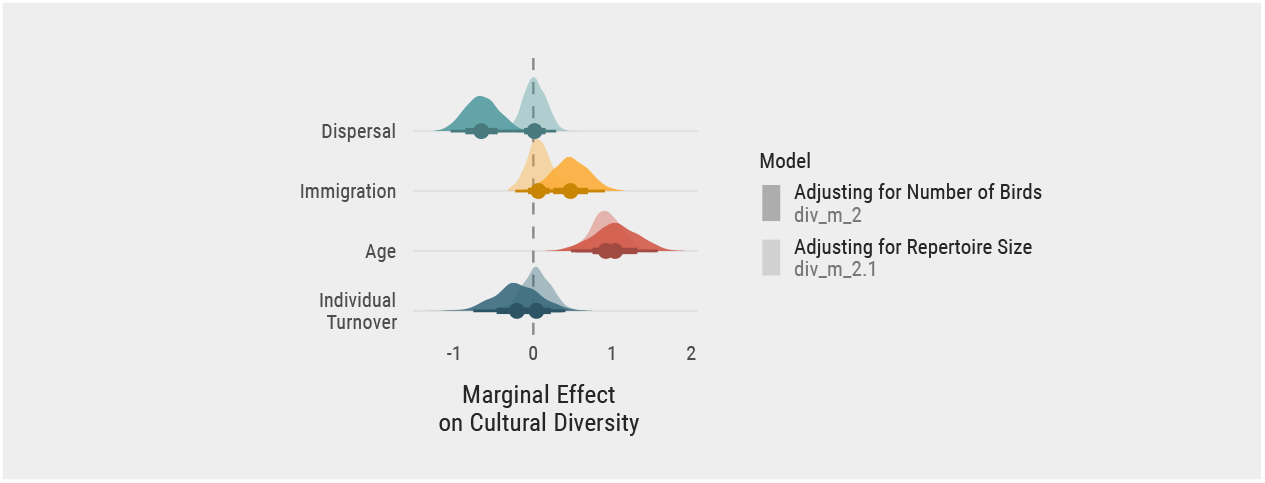
Marginal effects of demographic variables on absolute cultural diversity. Marginal effects of our predictor variables on absolute cultural diversity (the number of different song types sampled in a neighbourhood), while adjusting for the effect of either number of individuals (higher opacity fill, corresponding to model *div_m_2*) or number of song variants, including repeated variants (lower opacity fill, *div_m_2*.*1*).

**Figure S6.**
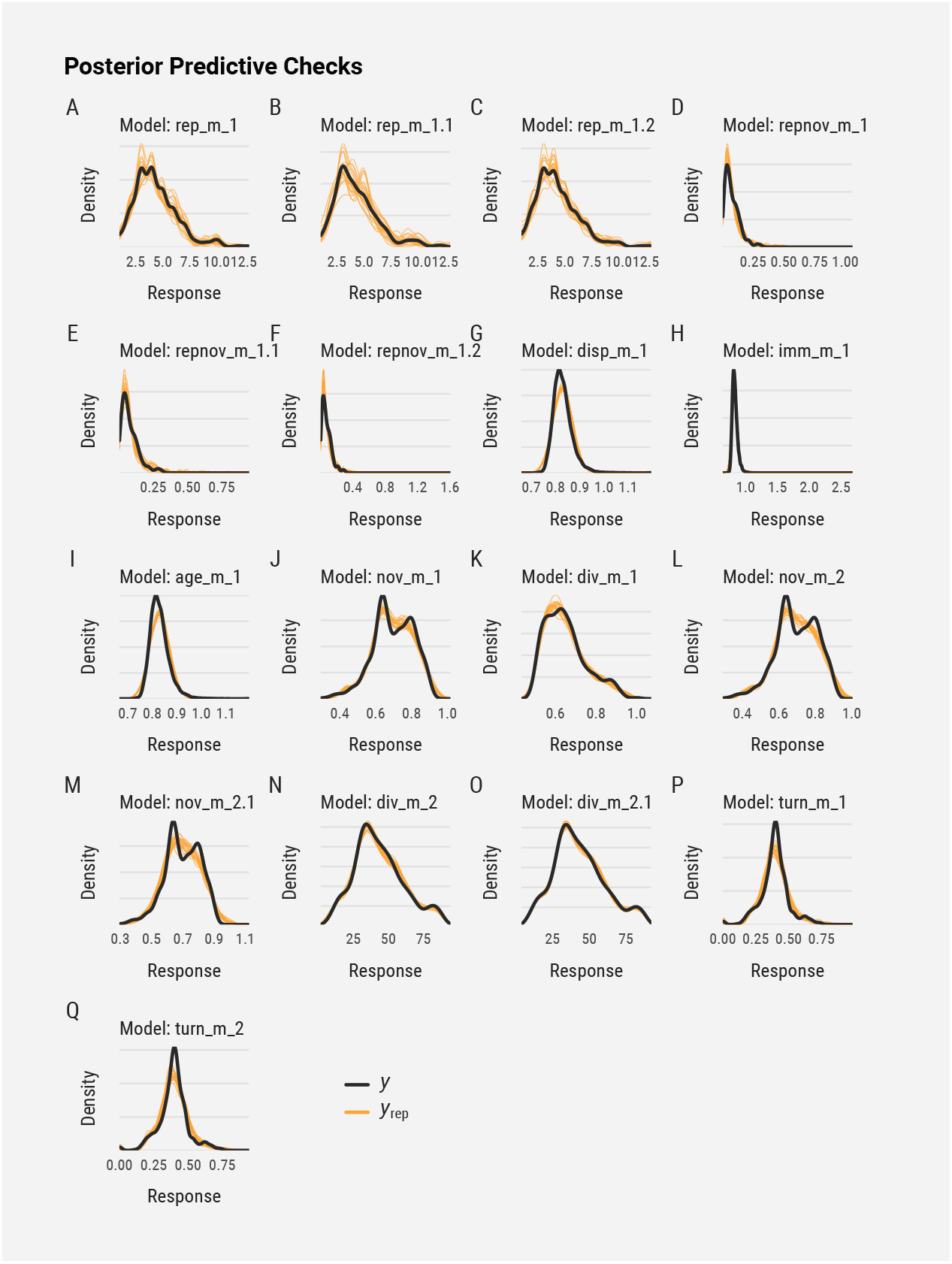
Posterior predictive checks for the main models in the study. Comparing simulations from the posterior predictive distribution *y*^*rep*^ (thin orange lines) with the outcome *y* (black lines) using Kernel density estimates. The posterior predictive distribution is a distribution of possible outcomes of the model given the data and the model parameters, here used to check the fit of the model to the data.

